# Classification of plant growth-promoting bacteria inoculation status and prediction of growth-related traits in tropical maize using hyperspectral image and genomic data

**DOI:** 10.1101/2022.03.04.483003

**Authors:** Rafael Massahiro Yassue, Giovanni Galli, Roberto Fritsche-Neto, Gota Morota

**Author notes:** Email addresses (RMY), (GG), (RFN), and (GM).

## Abstract

Recent technological advances in high-throughput phenotyping have created new opportunities for the prediction of complex traits. In particular, phenomic prediction using hyper-spectral reflectance could capture various signals that affect phenotypes genomic prediction might not explain. A total of 360 inbred maize lines with or without plant growth-promoting bacterial inoculation management under nitrogen stress were evaluated using 150 spectral wavelengths ranging from 386 to 1021 nm and 13,826 single-nucleotide polymorphisms. Six prediction models were explored to assess the predictive ability of hyperspectral and genomic data for inoculation status and plant growth-related traits. The best models for hyperspectral prediction were partial least squares and automated machine learning. The Bayesian ridge regression and BayesB were the best performers for genomic prediction. Overall, hyper-spectral prediction showed greater predictive ability for shoot dry mass and stalk diameter, whereas genomic prediction was better for plant height. The prediction models that simultaneously accommodated both hyperspectral and genomic data resulted in a predictive ability as high as that of phenomics or genomics alone. Our results highlight the usefulness of hyperspectral-based phenotyping for management and phenomic prediction studies.

**Core ideas:**

- Hyperspectral reflectance data can classify plant growth-promoting bacteria inoculation status
- Phenomic prediction performs better than genomic prediction depending on the target phenotype
- AutoML is a promising approach for automating hyperparameter tuning for classification and prediction

## Introduction

Addressing the growing food demand by increasing sustainable food production is critical in agriculture. The use of plant growth-promoting bacteria (PGPB) as inoculants to increase plant productivity and resilience to biotic and abiotic stresses has recently gained traction (Vejan et al., 2016; Majeed et al., 2018). However, an effective assessment of PGPB responses is difficult because the interaction between the PGPB × host genotype × environment is complex (Wintermans et al., 2016; Xiao et al., 2017). A method that can easily and accurately phenotype and predict plant growth-related traits under PGPB inoculation is needed (Rouphael et al., 2018; Susič et al., 2020).

Whole-genome molecular markers have been widely used for complex trait prediction (Meuwissen et al., 2001; Crossa et al., 2013; Fritsche-Neto et al., 2021). Genomic prediction performance mainly relies on the genetic relationship between individuals in the reference and target populations and linkage disequilibrium between genetic markers and quantitative trait loci. Prediction performance becomes suboptimal when the aforementioned relationship is weak or markers do not sufficiently capture quantitative trait loci signals (Habier et al., 2007; Windhausen et al., 2012; Sallam et al., 2015).

Hyperspectral cameras, sensors capable of capturing images in a wide spectrum of wavelengths, have recently been added to the realm of phenotyping tools available for plant genetics and breeding applications. It has been reported that the hyperspectral signatures of plant canopies are associated with plant nutrient status (Cilia et al., 2014; Mahajan et al., 2016), plant growth-related traits (Yang and Chen, 2004; Kaur et al., 2015), plant biomass (Jia et al., 2019; Ma et al., 2020), plant health (Lowe et al., 2017; Thomas et al., 2017), genotype discrimination (Chivasa et al., 2019), leaf water content (Ge et al., 2016), and soil microbial community composition (Carvalho et al., 2016). In particular, high-throughput phenotyping data-driven complex trait prediction, which is also known as phenomic prediction, is an active research topic for categorical and continuous phenotypes (Edlich-Muth et al., 2016; Rincent et al., 2018; Krause et al., 2019; Cuevas et al., 2019; Shu et al., 2021; Wang et al., 2021). Phenomic prediction is expected to capture the molecular composition of a plant, such as biochemical or physiological signals (endophenotypes), influencing phenotypes that genomic prediction may not directly explain (Rincent et al., 2018). Hyper-spectral reflectance data can be used to evaluate plant growth- or stress-related phenotypes in response to PGPB inoculation.

Several statistical learning models have been applied to phenomic prediction using hyperspectral image data. These include Bayesian ridge regression (BayesRR), least absolute shrinkage and selection operator (LASSO), partial least squares (PLS), BayesB, and neural networks (Montesinos-López et al., 2017b; Nigon et al., 2020; Qun’ou et al., 2021; Yoosefzadeh-Najafabadi et al., 2021). The use of the entire spectrum simultaneously, rather than selecting a small set of known wavelengths (e.g., spectra indices), can be beneficial for complex trait prediction (Aguate et al., 2017; Montesinos-López et al., 2017b,a). In general, fitting a machine learning model with good accuracy requires knowledge of the model structure to tune the hyperparameters. However, optimal hyperparameters are difficult to determine and often tuned manually using naive approaches (van Rijn and Hutter, 2018; Yang and Shami, 2020). As an alternative, automated machine learning (AutoML) has been proposed to reduce the need for human interference during the hyperparameter tuning process so that the application of machine learning becomes more automated and precise. (Feurer et al., 2015; Jin et al., 2019). Despite its potential, the use of AutoML for hyperspectral-based phenomic prediction of complex traits has not yet been fully explored. Therefore, the objectives of this study were to 1) evaluate the utility of hyperspectral image data to classify PGPB inoculation status, and 2) compare the predictive ability of genomic prediction, hyperspectral prediction, and their combination for growth-related phenotypes under PGPB management.

## Materials and Methods

### Plant growth-promoting bacteria experiment

A tropical maize association panel containing 360 inbred lines was used to study responses to PGPB. The inbred lines were evaluated with (B+) and without (B-) PGPB inoculation under nitrogen stress. The B+ management system consisted of a synthetic population of four PGPB, composed by *Bacillus thuringiensis* RZ2MS9, *Delftia* sp. RZ4MS18 (Batista et al., 2018, 2021), *Pantoea agglomerans* 33.1 (Quecine et al., 2012), and *Azospirillum brasilense* Ab-v5 (Hungria et al., 2010). The B-management included an inoculum with only liquid Luria-Bertani medium. Fertilization, irrigation, and other cultural practices were conducted according to the crop needs, except for nitrogen, which was not supplied. The phenotyping was performed when most of the plants had six expanded leaves. The manually measured phenotypes were plant height (PH), stalk diameter (SD), and shoot dry mass (SDM). Further information on the experimental design is available in Yassue et al. (2021a,b).

### Genomic data

A genotyping-by-sequencing method followed by the two-enzymes (PstI and MseI) protocol (Sim et al., 2012; Poland et al., 2012) was used to generate a total of 13,826 single-nucleotide polymorphisms (SNPs). The cetyltrimethylammonium bromide method was used to extract DNA (Doyle and Doyle, 1987). TASSEL 5.0 (Bradbury et al., 2007) was used to perform SNP calling with B73 (B73-RefGen v4) as the reference genome. The SNP markers were removed if the call rate was less than 90%, non-biallelic, or the minor allele frequency was less than 5%. Missing marker codes were imputed using Beagle 5.0 (Browning et al., 2018). Markers with pairwise linkage disequilibrium higher than 0.99 were removed using the SNPRelate R package (Zheng et al., 2012).

### Hyperspectral imaging and processing

Hyperspectral images of the maize lines grown under B+ and B – management conditions were obtained using a benchtop system of Pika L. camera (Resonon, Bozeman, MT, USA). The last expanded leaf was cut at the base of the stalk and immediately taken to the laboratory for imaging. The leaves were refrigerated using a cooler and gel refrigerant packs. Images were collected inside the dark room with a light supply to control for light variation. Radiometric calibration was performed using white and black tile panels approved by the camera system manufacturer. Each image cube (height, length, and band), consisted of 150 bands, varied from 386 to 1021 nm. The region of interest for each cube was the middle of the leaf. Once images were collected, the image processing step included applying a mask to remove the background and calculating the mean value of the reflectance from each pixel that referred to plant tissue. Hyperspectral image processing was performed using the Spectral Python (SPy) package. A summary of the data acquisition and processing is shown in Figure 1.

**Figure 1:**
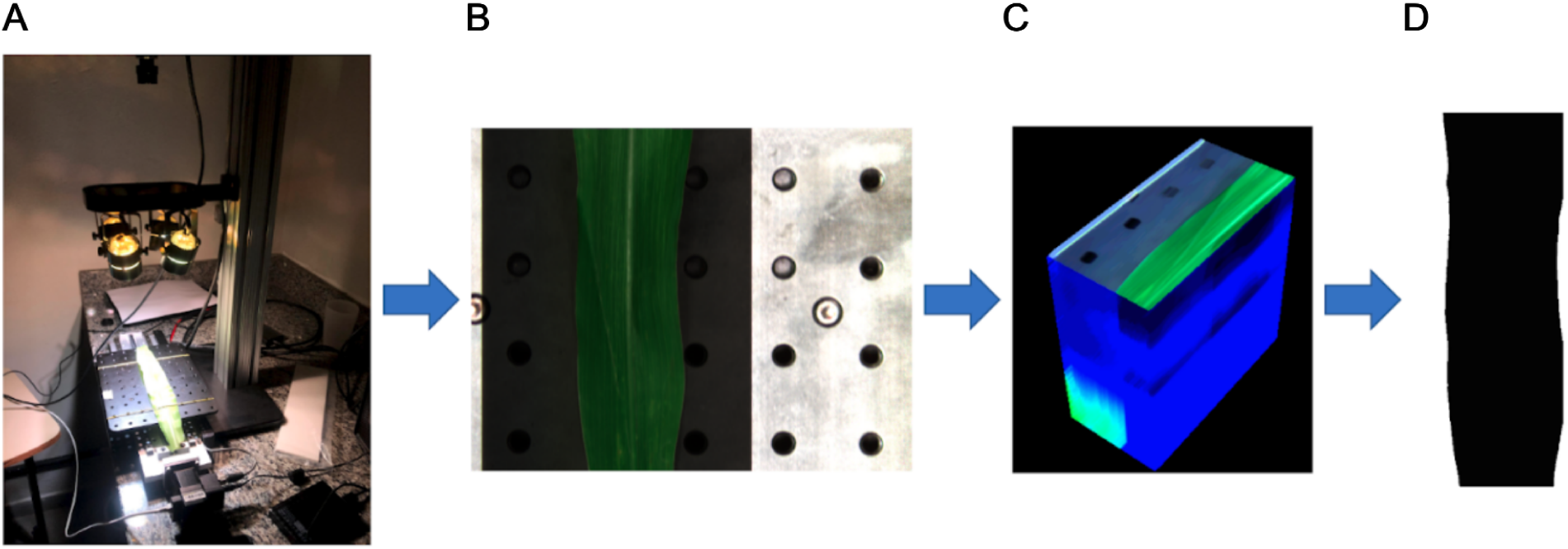
Summary of data acquisition and processing. A) the hyperspectral benchtop camera was used for data collection; B) region of interest of the last completed expanded leaf; C) cube image; and D) image mask.

### Prediction model

First, the utility of hyperspectral reflectance data to classify the PGPB inoculation status (B+ or B-) was evaluated. The purpose was to investigate whether the differences in genotypes’ biochemical or physiological signals between the two inoculation status are reflected at the hyperspectral level. If so, we expect to see a reasonable classification accuracy of the PGPB status using hyperspectral data. In this scenario, a classification accuracy of 0.5 is expected when genomic data are used as predictors because genomics is irrelevant to the presence or absence of inoculation. Second, the predictive abilities of genomic prediction, hyperspectral prediction, and their combination were compared for growth-related phenotypes, including PH, SD, and SDM.

The best linear unbiased estimators (BLUE) of genotypes were obtained using the following equation:

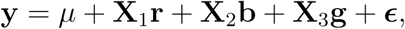

where **y** is the vector of phenotypes (PH, SD, SDM, and hyperspectral reflectance); **X**_1_, **X**_2_, and **X**_3_ are the incidence matrices for the fixed effects; *μ* is the intercept; **r, b**, and **g** are the fixed effects for replication, block within replication, and genotypes, respectively; and 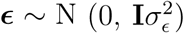 is the random residual effect, where **I** is the identity matrix and 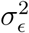 is the residual variance. The analysis was performed using the R package ASReml-R (Butler et al., 2017). The following classification and prediction models were used, which are summarized in Table 1.

**Table 1:** A list of models and covariates included in the analysis.

| Model | Classification | Prediction |  |  |
| --- | --- | --- | --- | --- |
|  | Hyperspectral image | Hyperspectral image | Genomics | Integration |
| LR<sup>1</sup> | ✓ |  |  |  |
| OLS<sup>2</sup> |  | ✓ |  |  |
| PLS-DA<sup>3</sup> | ✓ |  |  |  |
| PLS-R<sup>4</sup> |  | ✓ | ✓ |  |
| LASSO<sup>5</sup> | ✓ | ✓ | ✓ |  |
| BayesRR<sup>6</sup> | ✓ | ✓ | ✓ | ✓ |
| BayesB | ✓ | ✓ | ✓ | ✓ |
| AutoML<sup>7</sup> | ✓ | ✓ | ✓ |  |

<sup>1</sup> LR: logistic regresion
<sup>2</sup> OLS: ordinary least squares
<sup>3</sup> PLS-DA: partial least squares discriminant analysis
<sup>4</sup> PLS-R: partial least squares regression
<sup>5</sup> LASSO: least absolute shrinkage and selection operator
<sup>6</sup> BayesRR: Bayesian ridge regression
<sup>7</sup> Automated machine learning

#### Logistic regression and ordinary least squares

Logistic regression and ordinary least squares (OLS) were used to classify the inoculation status and predict growth-related traits, respectively, using the hyperspectral reflectance data. These models were not used for genomic prediction because the number of predictors was greater than the number of samples (maize lines).

#### Partial least squares

Partial least squares discriminant analysis (PLS-DA) and partial least squares regression (PLS-R) models (Wold et al., 2001) were fit using the caret R package (Kuhn, 2008). Partial least squares identifies latent variables that maximize the covariance between the predictors and responses while minimizing the error. The optimal number of the latent variables was estimated in the inner training set using k-fold cross-validation, with k set to four.

#### Least absolute shrinkage and selection operator

Least absolute shrinkage and selection operator was fit using the glmnet R package for linear regression as well as for generalized linear models (Friedman et al., 2010). The tuning parameter lambda was estimated by k-fold cross-validation with k set to four and was chosen according to the minimum mean cross-validation error.

#### BayesRR and BayesB

BayesRR and BayesB were fit for classification and prediction by treating the covariates as random (Meuwissen et al., 2001). For BayesRR, marker effects were sampled from a univariate normal distribution with a null mean and marker variance 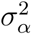, having a scaled inverse *χ*^2^ prior with scale parameter *S*_*α*_ and *ν*_*α*_ = 4 degrees of freedom. BayesB assumes that a priori, the marker effects have identical and independent mixture distributions, where each has a point mass at zero with a probability *π* and a scaled *t* distribution with a probability 1-*π* having a null mean and marker variance 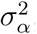, scale parameter *S*_*α*_, and *ν*_*α*_ = 4 degrees of freedom. The mixture parameter, *π*, was set to 0.99. For both BayesRR and BayesB, a flat prior was assigned to the intercept. The scale parameter was chosen such that the prior mean of 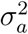 equals half of the phenotypic variance. All the Bayesian models were fitted using 60,000 Markov chain Monte Carlo samples, 6,000 burn-ins, and a thinning rate of 60 implemented in JWAS (Cheng et al., 2018). Model convergence was assessed using trace plots of the posterior means of the parameters.

#### Automated machine learning

Auto-sklearn automated machine learning aims to tune hyperparameters automatically by leveraging meta-learning, Bayesian optimization, and ensemble learning. The automated machine learning algorithm uses a meta-learning process that is quick but roughly explores the entire machine learning configuration space, which is then followed by Bayesian optimization to fine-tune the hyperparameters for performance. Finally, the ensemble process combines several machine learning models with different weights to increase predictive ability. The time limits for searching for an appropriate model and a single call were set to 900 s and 30 s, respectively. The AutoSklearnClassifier and AutoSklearnRegressor functions from AutoSklearn 0.14.2 (Feurer et al., 2015) were used for classification and prediction, respectively.

#### Multi-omic prediction

Multi-omic data integration may increase prediction performance relative to single-omic data (Krause et al., 2019; Galán et al., 2020; Guo et al., 2020). Multiomic prediction was performed by combining hyperspectral and genomic data using BayesRR and BayesB for growth-related phenotypes by setting different priors for each omic covariate. The framework closely followed that of Gonçalves et al. (2021) and Baba et al. (2021). The mixture parameter *π* was set to 0.99 for hyperspectral and genomic terms in multi-omics BayesB.

### Predictive performance

We used repeated random sub-sampling cross-validation to evaluate model performance. We split the population into training (80%) and test (20 %) sets, while maintaining balanced classes for inoculation categories (B+ and B-) and genotypes (Figure 2). The B+ and B-management conditions were jointly used for the classification models. Five genotypes were removed from the analysis to maintain a balanced structure because they were present only in one management. The training set was further split into inner training and validation sets for the models that required hyperparameter tuning. The inner training set was used for fine-tuning hyperparameters. The final model performance was evaluated in an independent test set that was not used in model training (Figure 2A). The accuracy of classification was derived using the following formula:

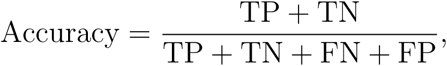

where TP, TN, FN, and FP are the number of genotypes in true positive, true negative, false negative, and false positive classes, respectively.

**Figure 2:**
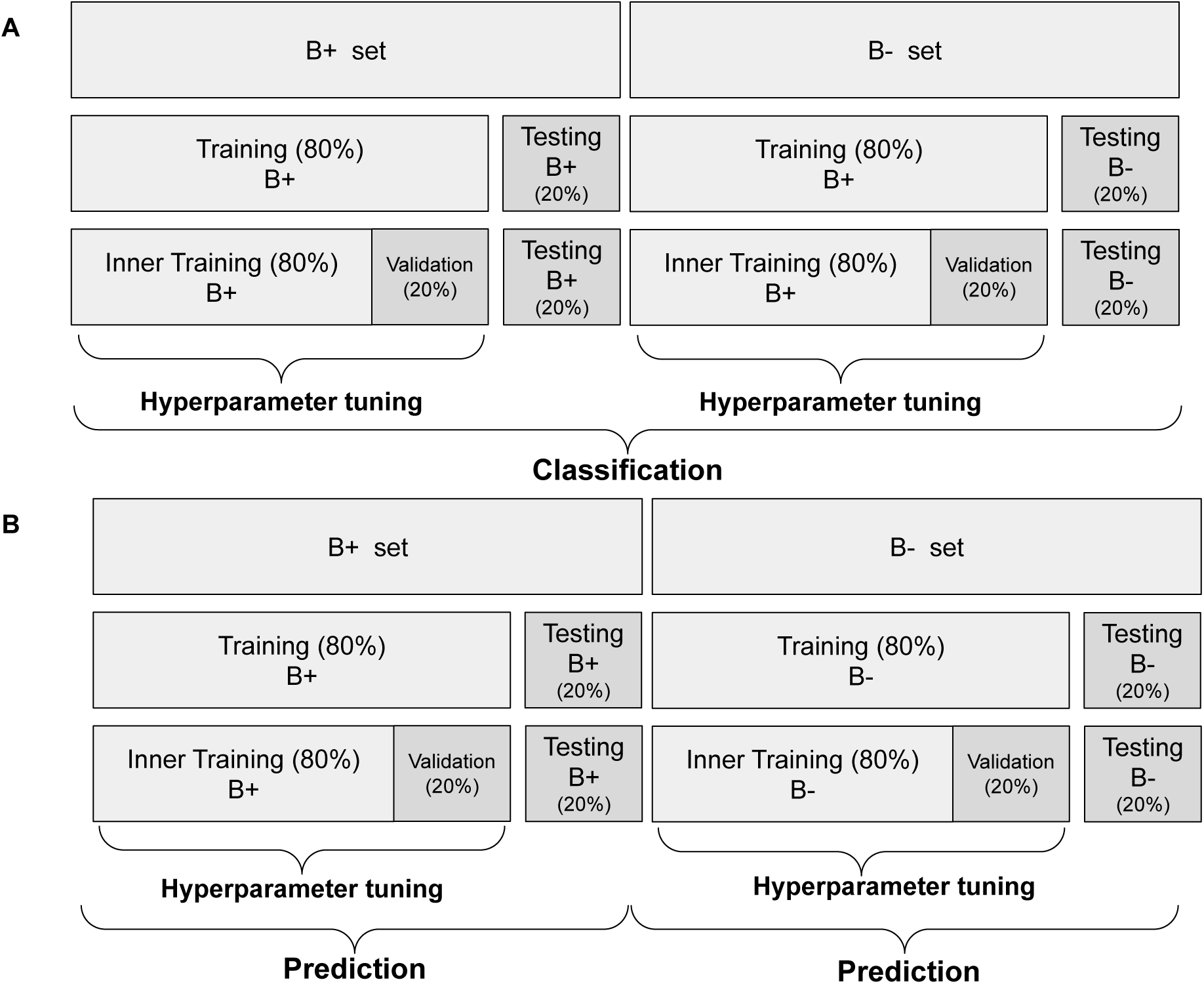
Graphic representation of a cross-validation design using the hyperspectral image and genomic data. The data were divided into training and testing sets. For PLS, LASSO, and Auto-sklearn, the training set was split into inner training and validation sets to perform hyperparameter tuning. This process was repeated 20 times using repeated random subsampling cross-validation. A) Classification was performed jointly using with (B+) and without (B-) plant growth-promoting bacteria inoculation conditions. B) Prediction was performed separately for each management condition.

For growth-related phenotypes, predictions were performed separately for each management condition (B+ and B-) (Figure 2B). The hyperparameters were tuned in the inner training set, similar to the classification tasks. We did not consider the interaction effect between the genotype and management condition because of the lack of such an effect (Yassue et al., 2021a,b). The predictive ability of the model was assessed using the Pearson correlation between the predictive values and BLUE of the genotypes.

## Results

### Correlation between growth-related phenotypes and hyperspectral bands

The hyperspectral signature and principal component biplot of the maize genotypes are shown in (Figure 3). No apparent visual patterns or clusters distinguished between the management conditions across the 150 hyperspectral bands. High hyperspectral variability was observed at the green and near-infrared wavelengths. The correlation matrix of the hyperspectral reflectance data showed that nearby bands had a higher correlation. As the distance between bands increased, the correlation decreased (Figure S1).

**Figure 3:**
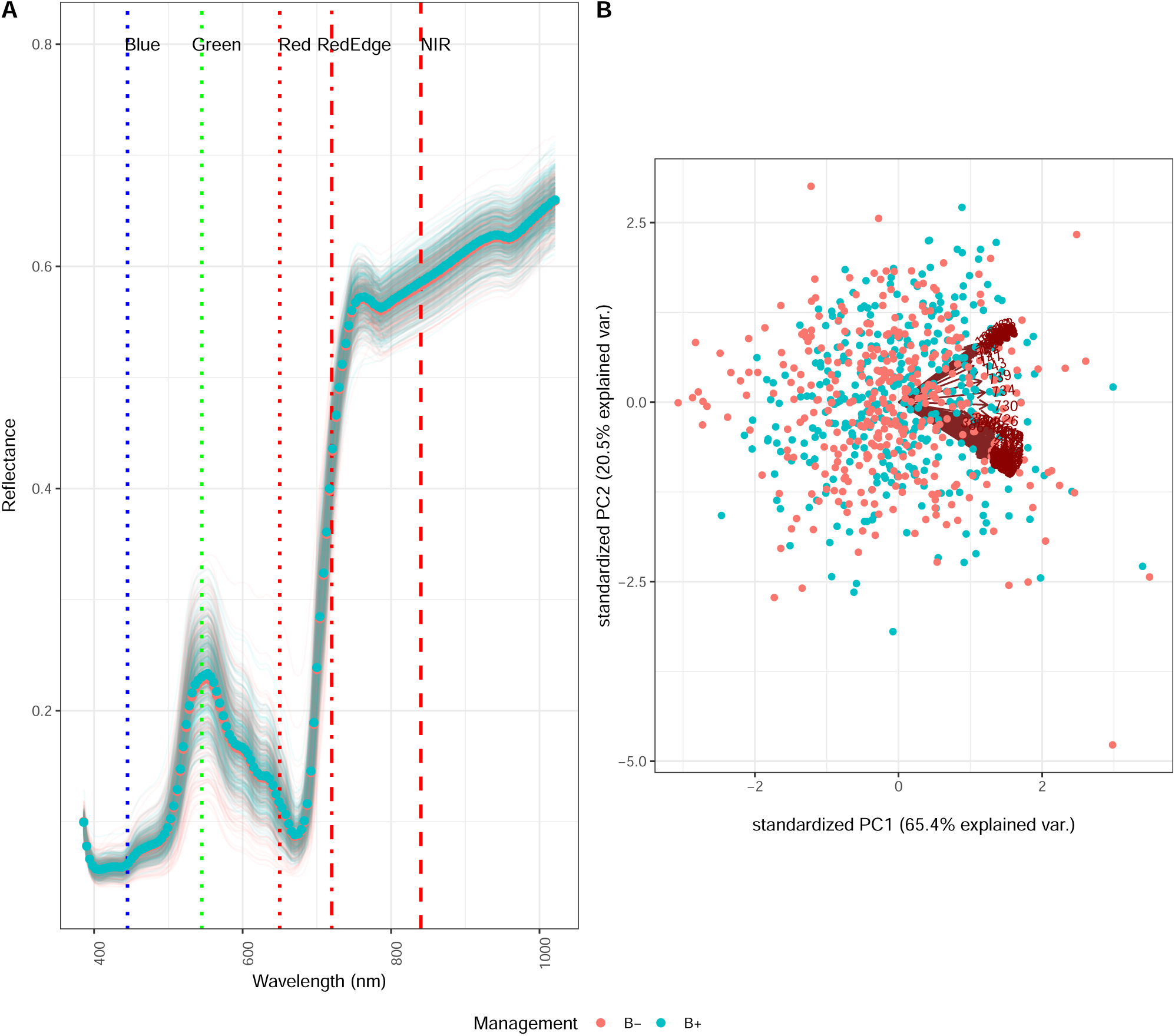
A: Spectral signatures of the maize genotypes under with (B+) and without (B-) plant growth-promoting bacteria mangement conditions. B: Principal component biplot of the 360 maize genotypes and 150 spectral wavelengths under with (B+) and without (B-) plant growth-promoting bacteria. Each point and arrow represent a genotype and a spectral wavelength, respectively.

The correlation between the single-band reflectance and growth-related phenotypes varied between -0.1 to 0.40, depending on the target phenotype and management condition (Figure 4). Overall, SDM was the most correlated trait with band reflectance values. Additionally, the correlation with band reflectance was higher for the inoculated samples (B+). Most of the bands were associated with the target phenotypes and the correlation peaks occurred between blue and green for PM and between RedEdge and near-infrared for SD and SDM.

**Figure 4:**
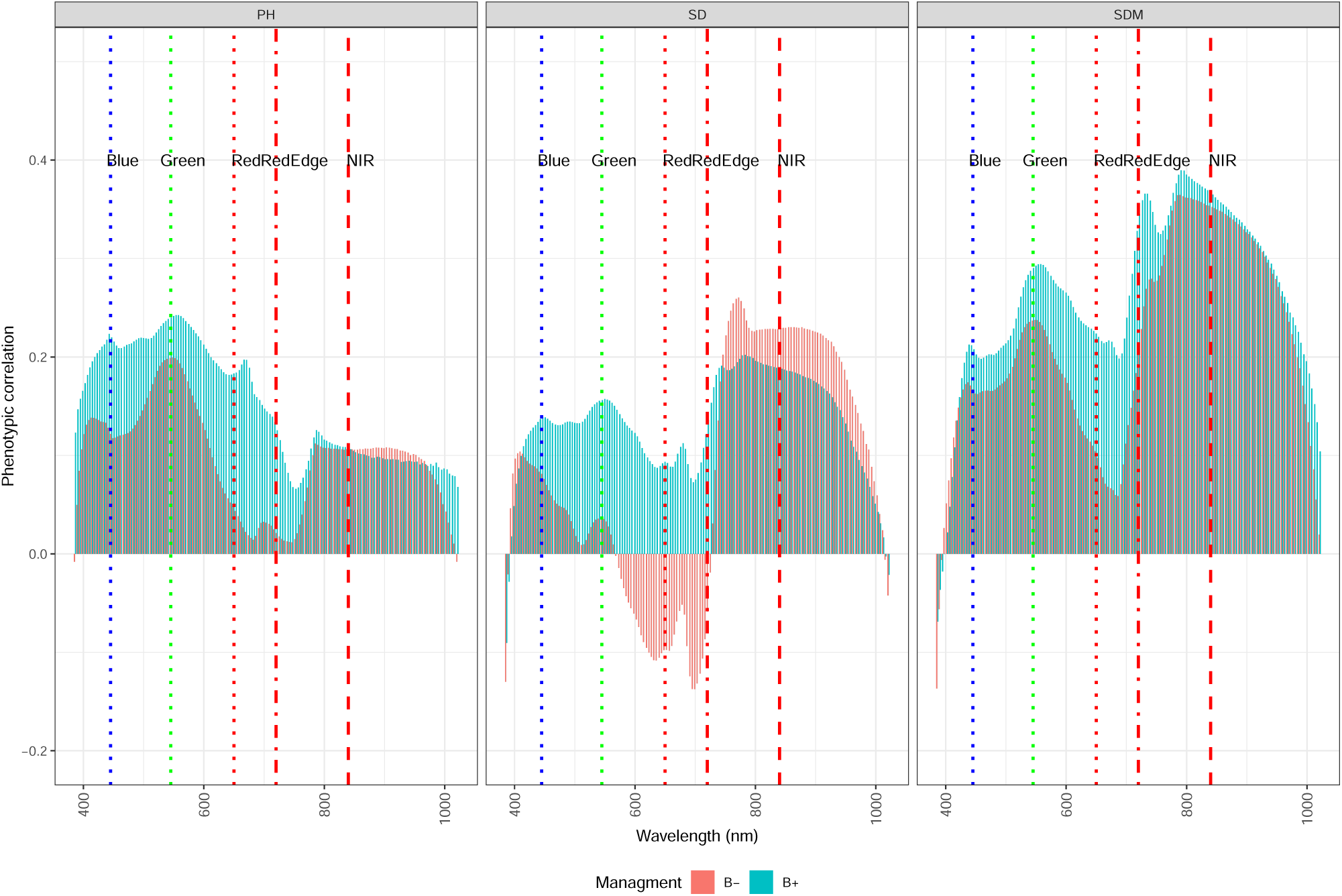
Correlations between manually measured growth-related phenotypes and hyper-spectral reflectance values under the two management conditions (B+ and B-). PH: plant height; SD: stalk diameter; and SDM: shoot dry mass.

### Classification of inoculation status

The accuracy of classifying the inoculation status (B+ and B-) using hyperspectral reflectance data is shown in Figure 5. AutoML and PLS were the best classification models, followed by OLS and LASSO. The accuracy achieved by AutoML and PLS was higher than 0.8, demonstrating that the hyperspectral profiles of the B + and B-inoculation status were distinct. However, BayesRR and BayesB did not perform well, and their performance was worse than that of OLS.

**Figure 5:**
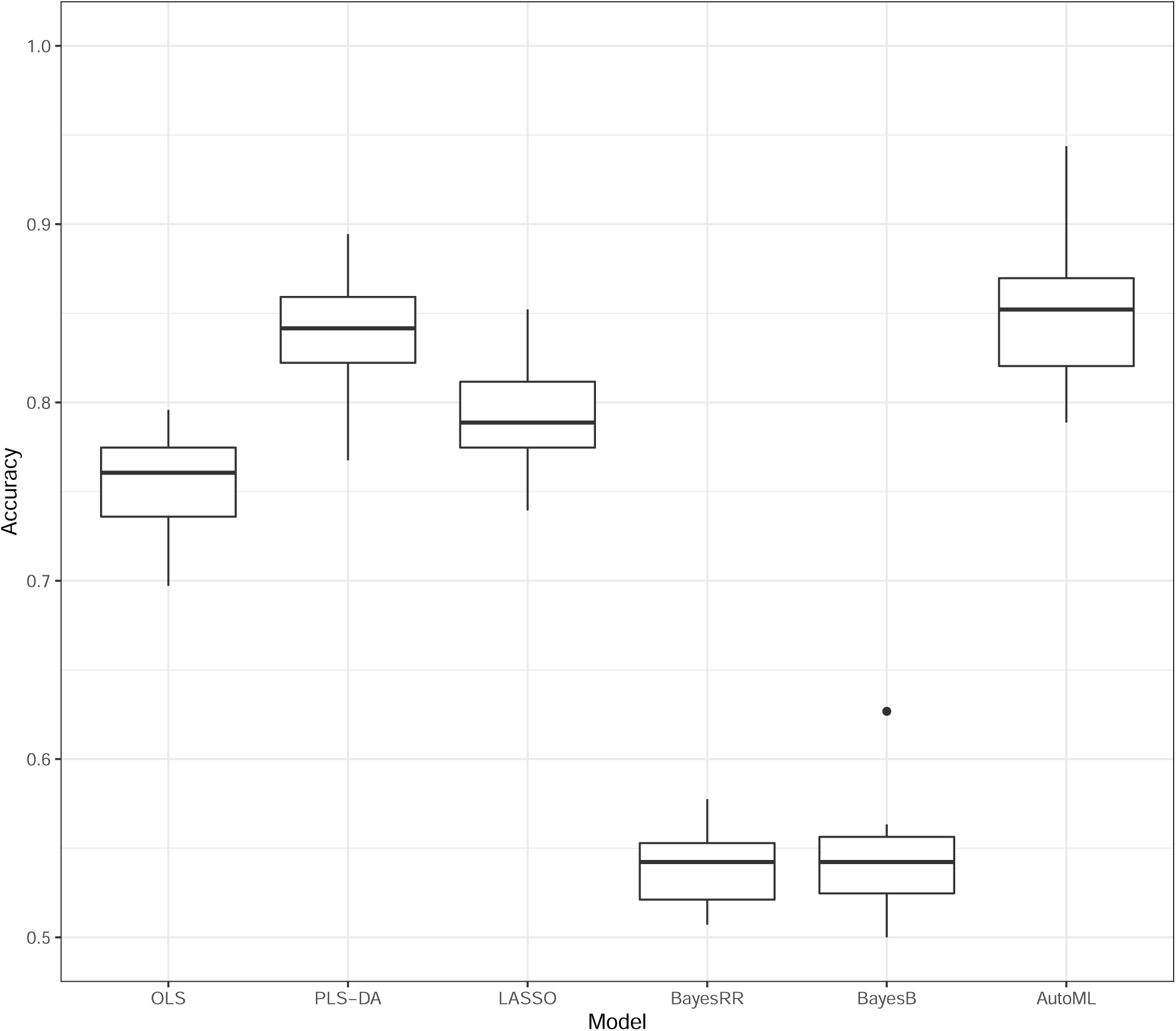
Classification accuracy of inoculation status (B+ and B-) using 150 hyperspectral bands. OLS: ordinary least squares; PLS-DA: partial least squares discriminant analysis; LASSO: least absolute shrinkage and selection operator; BayesRR: Bayesian ridge regression; and AutoML: automated machine learning.

### Prediction of growth-related phenotypes

The performance of hyperspecral-driven phenomic prediction and SNP-driven genomic prediction is shown in Figure 6. We obtained the highest and lowest predictions for PH and SDM, respectively, using phenotypic prediction. Furthermore, we observed higher predictions for B+ than for B- in PH, whereas the opposite was observed for SD. The predictions were higher for B+, except when BayesB was used. AutoML, PLS, and LASSO performed equally well in predicting SD and SDM. In contrast, no notable differences were observed in PH. In genomic prediction, the best prediction was obtained for PH. The B+ management conditions were more predictable than B-. Overall, the predictive performance was not sensitive to the choice of the genomic prediction model. The multi-omics models improved the prediction correlations for all phenotypes compared with the single omics model; however, this gain was incremental.

**Figure 6:**
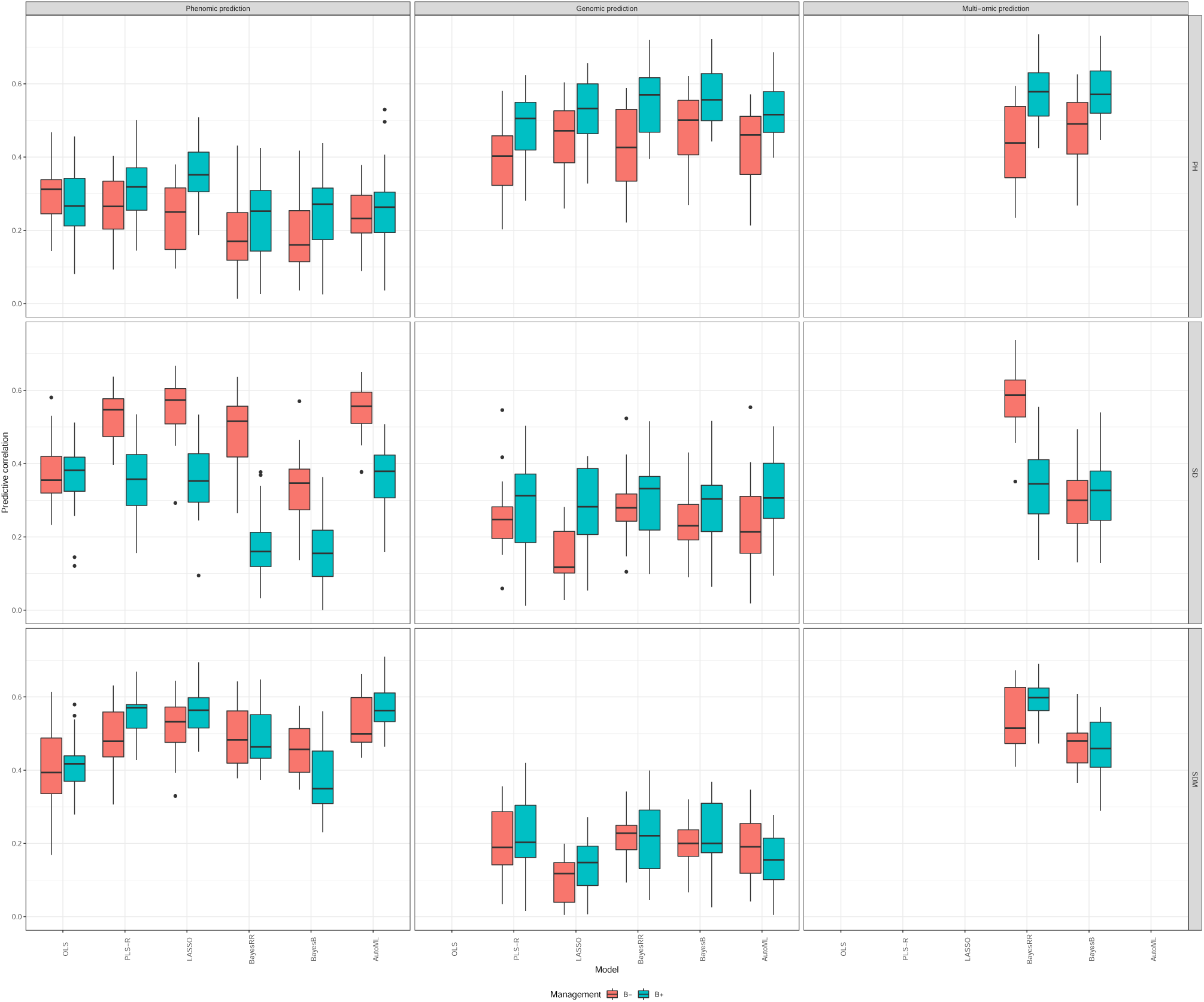
Predictive ability of plant height (PH), stalk diameter (SD), and shoot dry mass (SDM) using phenomic prediction, genomic prediction, and multi-omic prediction models in each management condition (B+ and B-). OLS: ordinary least squares; PLS-R: partial least squares regression; LASSO: least absolute shrinkage and selection operator; BayesRR: Bayesian ridge regression; and AutoML: automated machine learning.

## Discussion

### Phenotyping PGPB response

Recent studies have found that inoculation with PGPB can modify plant structure and increase plant biomass and resilience to nitrogen stress (Yassue et al., 2021a,b). This study evaluated PGPB responses at the level of hyperspectral reflectance data. The ability to classify the inoculation status using phenomics showed that the hyperspectral camera could capture signals unique to each management condition. Previous studies have reported that the PGPB species used in this study are capable of producing indole acetic acid, fixing nitrogen, and promoting phosphate solubilization (Quecine et al., 2012; Batista et al., 2018, 2021). A field trial study reported that *Azospirillum brasilense* can increase nitrogen, potassium, boron in the leaves of maize (Hungria et al., 2010). However, indole acetic acid production or nutrient status was not evaluated in the present study, and the interpretation of the hyperspectral signature is limited. The results of our study align with those of Carvalho et al. (2016), who showed that leaf hyperspectral patterns in winter wheat have the potential to detect changes in soil microbial communities. Moreover, Susič et al. (2020) reported successful classification of PGPB inoculation status for nematicidal effects in tomatoes using hyperspectral image data. They found that the hyperspectral signature can be used to assess plant stress after inoculation with PGPB. The minor difference in hyperspectral reflectance curves between the B+ and B-management conditions, in addition to the lack of clustering in PCA, suggests that most of the bands probably contributed to inoculation status classification. Further studies should be conducted to evaluate the structural, morphological, and chemical differences between B+ and B-management conditions.

### Model performance

The statistical modeling of high-throughput phenotyping data in quantitative genetics is becoming increasingly important (Morota et al., 2022). We evaluated the utility of hyperspectral-based phenomic prediction to classify PGPB inoculation status and compared the predictive ability of genomic prediction, hyperspectral prediction, and data integration for growth-related phenotypes using statistical prediction models in tropical maize. Generally, PLS and AutoML were competitive in many scenarios and the performance of PLS in our study agreed with previous work that used smaller datasets (Fu et al., 2019; Galli et al., 2020; Shu et al., 2021). Montesinos-López et al. (2017b) also reported that PLS-R performed better than BayesB for predicting wheat yield using hyperspectral reflectance.

AutoML performed equally well or better than PLS for phenomic classification, suggesting the usefulness of hyperparameter tuning and ensemble learning in machine learning (Figure 5). We obtained better predictions in the B+ management conditions for PH, SDM, and for SD in the B-management conditions. This could be explained by the extent of correlation between growth-related phenotypes and hyperspectral reflectance (Figure 3).

For genomic prediction, the higher predictive ability for PH was probably due to its higher heritability in comparison to SD and SDM (Yassue et al., 2021a,b). Similar to the phenomic prediction, AutoML yielded relatively good predictions. In addition to AutoML, BayesRR and BayesB were competitive and stable across the three phenotypes investigated. This was expected because these models are well accepted in the genomic prediction literature. However, BayesRR and BayesB did not perform well with hyperspectral reflectance data. Simple OLS outperformed BayesRR and BayesB in some cases, suggesting that shrinkage or variable selection is not necessarily beneficial when the number of predictors is lower than the number of samples (Whittaker et al., 2000). The strength of OLS is that the estimated effect has the property of the best linear unbiased estimator, and its expectation is equal to the true effect if the Gauss–Markov theorem is satisfied (Searle and Gruber, 2016). This property appears to be useful for hyperspectral prediction.

### Phenomic vs genomic prediction

The hyperspectral signature of plants is considered an endophenotype because it can capture the expression of genotypes under specific conditions and is useful in predicting complex traits (Rincent et al., 2018). We found that phenomic prediction was more predictive than genomic prediction for SD and SDM. In contrast, genomic prediction was better than phenomic prediction for PH. Although the multi-omic model produced the highest predictive correlations for all three phenotypes by integrating genomic and hyperspectral information, there was no noticeable enhancement over the best single-omic prediction. Our results are consistent with those of previous studies (Xu et al., 2017; Goncalves et al., 2021) reporting for different species or omic data combinations. Overall, our results showed that the hyper-spectral signature of genotypes is a valuable resource for complex trait prediction. Further studies are needed to improve hyperspectral image acquisition for greenhouse experiments.

The genetic gain equation, also known as the breeder’s equation, is defined as 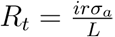, where *R*_*t*_ is the response to selection by time, *i* is the selection intensity, *r* is the selection accuracy, *σ*_*a*_ is the square root of additive genetic variation, and *L* is the generation interval (Cobb et al., 2019). Phenomic prediction has the potential to alter *i* and *r* because it can increase the selection intensity and accuracy by phenotyping a larger number of genotypes. Conversely, genomic prediction may also increase accuracy and reduce the generation interval. Because the hyperspectral signature is an endophenotype, it can be influenced by environmental effects, unlike genomic information, which is specific to the individual. Hence, the use of phenomic or genomic prediction models depends on the goal of selection (Hickey et al., 2017).

The disadvantage of using a benchtop camera is the need to bring plants to an imaging room. In this study, the region of interest was the middle portion of the leaf, which required manual collection of maize leaves. This could limit the applicability hyperspectral imaging in plant breeding or genetics program pipelines because of the laborious data collection time. The use of hyperspectral cameras in a low-cost, high-throughput phenotyping platform, such as that reported by Yassue et al. (2021b), may ease the application of phenomic prediction that uses hyperspectral reflectance data.

## Conclusions

We found that hyperspectral reflectance data were useful predictors for classifying PGPB inoculation status and predicting growth-related phenotypes in tropical maize. Phenomic prediction showed better performance than genomic prediction for SD and SDM. AutoML is a promising approach for classification and prediction tasks that mitigate manual hyper-parameter tuning. The integration of hyperspectral and genomic data resulted in predictive performance as high as that of the best single-omic model. Overall, our results highlight the usefulness of hyperspectral imaging in management and phenomic prediction studies.

## Acknowledgments

The authors acknowledge Pedro Takao Yamamoto and Fernando Henrique Iost Filho for their support in collecting the hyperspectral images.

## Author contributions

Rafael Massahiro Yassue: Conceptualization; Data curation, Formal analysis, Investigation; Methodology; Visualization; Writing-original draft; Writing-review & editing. Giovanni Galli: Investigation; Methodology; Writing-review & editing. Roberto Fritsche-Neto: Conceptualization; Funding acquisition; Supervision; Writing-review & editing. Gota Morota: Conceptualization; Methodology; Funding acquisition; Supervision; Writing-original draft; Writing-review & editing.

## Abbreviations

(AutoML): automated machine learning
(BayesRR): Bayesian ridge regression
(BLUE): best linear unbiased estimators
(LASSO): least absolute shrinkage and selection operator
(OLS): ordinary least squares
(PLS): partial least squares
(PLS-DA): partial least squares discriminant analysis
(PLS-R): partial least squares regression
(PGPB): plant growth-promoting bacteria
(PH): plant height
(SDM): shoot dry mass
(SNP): single-nucleotide polymorphism
(SD): stalk diameter
(B+): with plant growth-promoting bacteria inoculation
(B-): and without plant growth-promoting bacteria inoculation

