## Supplementary material for "Classification of plant growth-promoting bacteria inoculation status and prediction of growth-related traits in tropical maize using hyperspectral image and genomic data": Figure S1

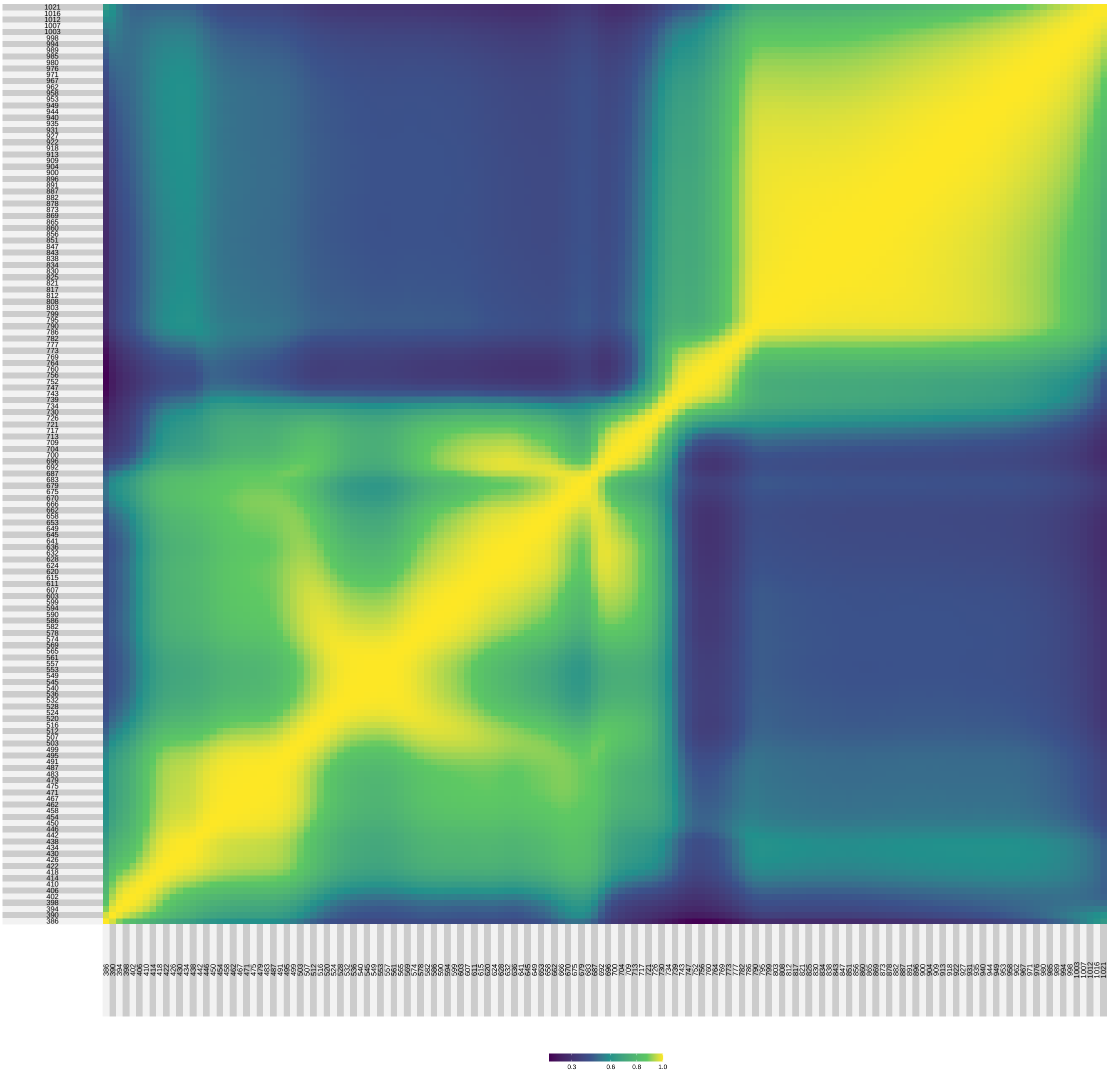


Figure S1: Correlation matrix of the hyperspectral data ranging from 386 to 1021 nm. The correlation coefficients were computed from the averages of the two management conditions.
